# *In vitro* toxicity and efficacy of verdinexor, an exportin 1 inhibitor, on opportunistic viruses affecting immunocompromised individuals

**DOI:** 10.1101/351825

**Authors:** Douglas G. Widman, Savanna Gornisiewicz, Sharon Shacham, Sharon Tamir

## Abstract

Infection of immunocompromised individuals with normally benign opportunistic viruses is a major health burden globally. Infections with viruses such as Epstein-Barr virus (EBV), human cytomegalovirus (HCMV), Kaposi’s sarcoma virus (KSHV), adenoviruses (AdV), BK virus (BKPyV), John Cunningham virus (JCPyV), and human papillomavirus (HPV) are significant concerns for the immunocompromised, including when these viruses exist as a co-infection with human immunodeficiency virus (HIV). These viral infections are more complicated in patients with a weakened immune system, and often manifest as malignancies resulting in significant morbidity and mortality. Vaccination is not an attractive option for these immune compromised individuals due to defects in their adaptive immune response. Verdinexor is part of a novel class of small molecules known as SINE (Selective Inhibitor of Nuclear Export) compounds. These small molecules demonstrate specificity for the nuclear export protein XPO1, to which they bind and block function, resulting in sequestration of XPO1-dependent proteins in the nucleus of the cell. In antiviral screening, verdinexor demonstrated varying levels of efficacy against all of the aforementioned viruses including previously with HIV. Studies by other labs have discussed likely mechanisms of action for verdinexor (ie. XPO-1-dependence) against each virus. GLP toxicology studies suggest that anti-viral activity can be achieved at a tolerable dose range, based on the safety profile of a previous phase 1 clinical trial of verdinexor in healthy human volunteers. Taken together, these results indicate verdinexor has the potential to be a broad spectrum antiviral for immunocompromised subjects for which vaccination is a poor option.

## Introduction

Immunocompromised patients demonstrate susceptibility to latent opportunistic infections, particularly when concurrently infected with HIV. There are a number of dsDNA viruses that threaten those with immune deficiencies (Table 1). Vaccines are likely to suffer from waning CD4^+^ T cell counts and thus antiviral therapies are a promising therapeutic stragegy. Despite the prevalence of opportunistic viral infections worldwide, there are few treatments and vaccines available for those described here. Thus, there is an unmet need for novel treatments.

**Table 1.** Charecterstics of HIV-associated severe viral infections

| <u>Virus</u> | <u>Virus Type</u> | <u>Genome</u> | <u>Virus Family</u> | <u>Transmission</u> | <u>Diseases</u> | <u>Prevalence</u> |
| --- | --- | --- | --- | --- | --- | --- |
| **Epstein-Barr virus (EBV)** | Enveloped, dsDNA | 184 kb | Herpesviridae | Spread by saliva and sexual fluids | • Infectious mononucleosis<br>• oral hairy leukoplakia<br>• Burkitt's lymphoma<br>• Hodgkin's lymphoma<br>• autoimmune diseases | • 200,000 cancer cases per year attributed to EBV<br>• 90% of population has evidence of previous infection<br>• 143,000 deaths in 2010 attributed to EBV |
| **Human Cytomegalovirus (HCMV)** | Enveloped, dsDNA | 220 kb | Herpesviridae | Spread by body fluids (urine, saliva, sexual fluids) | • mononucleosis, glandular fever, pneumonia | • Infects between 60-70% of adults in industrialized countries and almost 100% in emerging countries |
| **Kaposi's Sarcoma-associated Herpesvirus (KSHV)** | Enveloped, dsDNA | 165 kb | Herpesviridae | Sexually transmitted | • cancer in HIV-infected individuals | • Major variations in prevalence geographically<br>• Seropositivity rates >50% in sub-Saharan Africa, 20-30% in Mediterranean Countries, <10% in most of Europe, Asia and US<br>• Prevalence is elevated in men who have sex with men |
| **BK Virus** | Non-enveloped, dsDNA | 5 kb | Polyomaviridae | Spread by body fluids (urine, saliva); fecal-oral | • neuropathy and malignancies including brain tumors, osteogenic sarcomas, urinary tract neoplasms, meningiomas | • More common in young children; seroprevalence rates of 65-90% being reached by the age of 10 years old |
| **John Cunningham Virus (JCV)** | Non-enveloped, dsDNA | 5 kb | Polyomaviridae | Spread by body fluids (urine, saliva); fecal-oral | • leukemias<br>• lymphomas<br>• reactivation of viral infection that can lead to progressive multifocal leukoencephalopathy (PML) | • JCV infection prevalence varies between populations<br>• Adult JCV prevalence rates between 20-60% (increases with age) |
| **Human Papilloma Virus (HPV)** | Non-enveloped, dsDNA virus with a circular genome | 8 kb | Papillomaviridae | Sexually transmitted | • cervical cancer<br>• genital warts<br>• cancers of the vulva, vagina, penis and anus | • 79 million Americans currently infected (14 million new cases annually) |
| **Adenovirus (AdV)** | Non envelope | 36 kb | Adenoviridae | Airborne transmission as well as via direct and indirect contact with infected persons | • Various respiratory illnesses<br>• Cold like symptoms: sore throat, bronchitis, pneumonia, diarrhea and conjunctivitis<br>• Acute respiratory disease<br>• Multi-organ disease in immunocompromised population | • Exact prevalence and incidence is unknown<br>• Very common infection-responsible for 2-5% of all respiratory infections |

Exportin 1 (XPO1) mediates export of proteins containing leucine rich nuclear export signals (NES) and RNA transcripts [1–4]. Nucleocytoplasmic transport occurs through the nuclear pore complex, which allows passive diffusion of small molecules, but requires active transport of larger cargos. Nucleocytoplasmic trafficking pathways are integral to inflammation and the pathogenesis of numerous viral infections, where key aspects of the viral life cycle are dependent on XPO1-mediated transport. [5].

Verdinexor (KPT-335; Fig. 1), an orally bioavailable Selective Inhibitor of Nuclear Export (SINE) functions by covalently binding to the active site of XPO1, preventing docking of cargo molecules. Inhibition of XPO1 with verdinexor disrupts many viral life cycles by blocking the nuclear export of vRNP [6] and other critical viral components [7]. XPO1 inhibition also leads to nuclear retention of host factors essential to viral replication, and inflammatory molecules like cytokine transcripts and transcription factors responsible for immunopathology. Verdinexor attenuates inflammation by forcing nuclear retention of proinflammatory cytokine transcriptional inhibitors (such as IκB, RXRα, and PPARγ) (Fig. 2). XPO1-mediated translocation of the tumor suppressor protein GLTSCR2 to the cytoplasm results in attenuation of RIG-I and decreased interferon-β production [8]. Therefore, verdinexor has potential for treating viruses by inhibiting viral replication and relieving symptoms through suppression of inflammatory responses.

**Figure 1.**
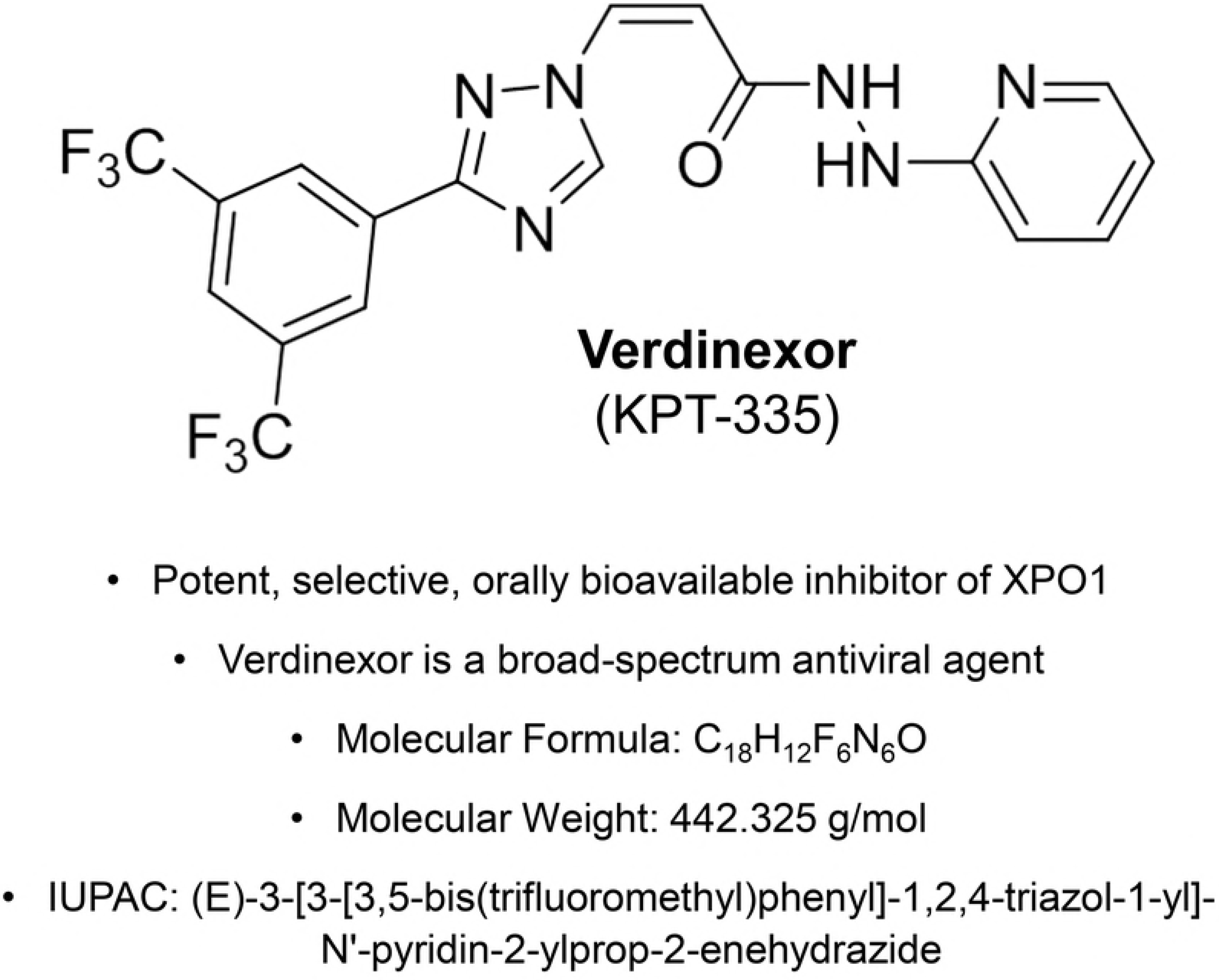
Chemical properties of verdinexor (KPT-335).

**Figure 2.**
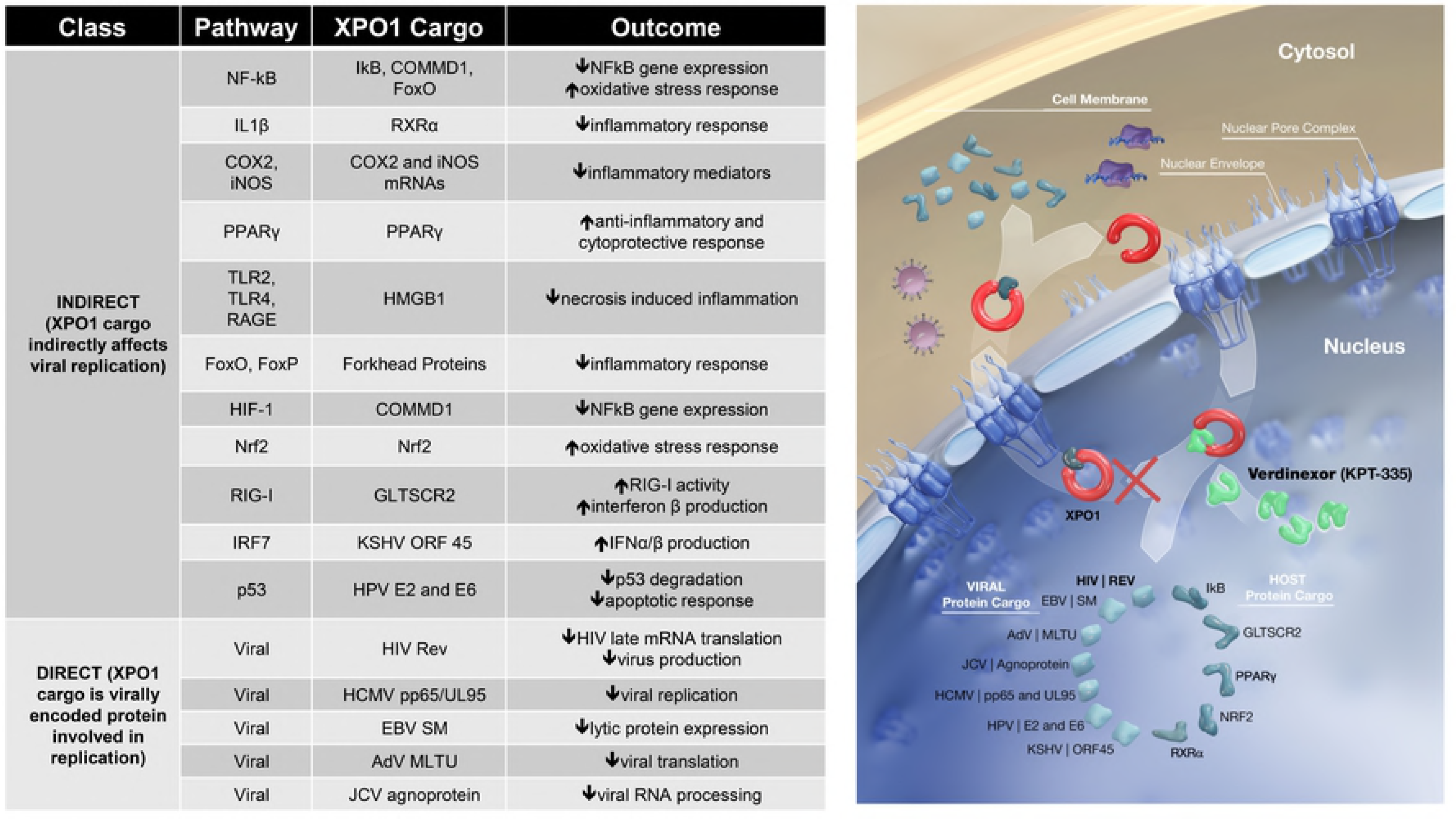
Critical XPO1 Cargoes. Molecules that rely on XPO1 for nuclear export are highlighted, including those that have either direct or indirect effects on viral replication and inflammation. This list is not exhaustive but represents cargoes that are important for viral life cycles.

To assess potential off-target toxicities associated with verdinexor, a panel of in vitro protein binding assays were performed to evaluate the potential interaction of verdinexor with 104 principal receptor/ligand interactions (peptides, growth factors, ion channels, transporters, kinases, cysteine proteases (including caspases and matrix metalloproteinases) (Study No. KS-1016 and PE-50946871). At a concentration of 10 μM, none of the enzymes, receptors, transporters, kinases, or cysteine proteases were significantly affected by verdinexor treatment, with the exception of marginal activity for histamine H1 receptor binding (33% inhibition). Functional activity assays (antagonist, agonist) for histamine H1 receptor binding demonstrated that verdinexor IC50 values were >10 µM in both assays (Study No. KS-1017). A Phase 1, Randomized, Double-Blind, Placebo-Controlled, Sequential, Dose-Escalating Trial to Evaluate the Safety and Tolerability of Oral Verdinexor (KPT-335) in Healthy Adult Subjects was conducted in Nucleus Network, Melbourne, Victoria, Australia [ClinicalTrials.gov: NCT02431364]. No study results are currently posted for this study. Furthermore, a clinical study was submitted to the USA FDA on IND#122718. This phase 1a, Randomized, Double-Blind, Placebo-Controlled, Sequential, Dose-Escalating Trial of Single and Multi-Dose oral verdinexor (KPT-335) to Evaluate the Safety and Tolerability in Healthy Elderly Volunteers is yet to be initiated and recruit subjects.

XPO1 mediates the nuclear export of the HIV *Rev* protein and we have shown that treatment with verdinexor and other SINE compounds results in forced nuclear retention of Rev-GFP fusion at an effective concentration (EC_50_) of 160nM [7], and anti-HIV activity in PBMCs from healthy donors at an EC_50_±116 ± 54nM [7]. Reduced expression of late viral mRNA was also observed. SINE compounds exhibit significant *in vitro* antiviral efficacy against seven strains of HIV, indicating that verdinexor is active against the *Rev* proteins of diverse viral strains. Thus, there is rationale for inhibiting XPO1 as a treatment for HIV [7], and perhaps simultaneously the comorbidities that often manifest with infection.

Here, we demonstrate the antiviral properties of verdinexor against opportunistic dsDNA viruses. These data, along with our previous HIV observations, suggest that verdinexor may be able to safely inhibit multiple concurrent viral infections commonly associated with pathology in immunocompromised individuals. As such, we believe these data justify the evaluation of verdinexor in small animal models of viral diseases that demonstrated the highest sensitivity to verdinexor in this screen.

## Experimental Procedures

### Compound Characterization

Verdinexor (KPT-335; Fig. 1) was determined to be 99.7% pure by use of infrared spectroscopy and HPLC chromatography.

### Cells and viruses

EBV Akata strain was assayed in Akata cells (John Sixbey, LSU) latently infected with EBV (and therefore oncogenic). Cells were maintained in RPMI 1640, (Mediatech) with 10% FBS (Hyclone), L-glutamine, penicillin and gentamicin.

Human CMV strain AD169 was obtained from ATCC and assayed in human foreskin fibroblast cells (HFF). HFF cells were prepared at the University of Alabama at Birmingham (UAB) tissue procurement facility with IRB approval as previously described [9, 10].

Murine CMV strain Smith was obtained from Earl Kern and assayed in mouse embryonic fibroblast cells (MEF) immortalized by telomerization. Guinea pig CMV strain 22122 was obtained from the ATCC and assayed in primary guinea pig lung cells (GPL). Both MEF and GPL cells were prepared at the UAB tissue procurement facility with IRB approval as previously described [9, 10].

KSHV strain BCBL-1 was obtained as latently infected BCBL-1 cells through the NIH AIDS Research and Reference Reagent Program. BCBL-1 cells were maintained in growth medium consisting of RPMI 1640 supplemented with 10% FBS, penicillin, gentamicin, and l-glutamine.

Human adenovirus type 5 was obtained from ATCC and cultured in HeLa cells under standard conditions.

John Cunningham polyomavirus (JCPyV) strains MAD-1 and MAD-4 and BK polyomavirus (BKPyV) Gardner strain were obtained from ATCC. COS-7 cells were obtained from ATCC and maintained according to ATCC-supplied protocols.

HPV-11 and −18 were tested in HEK293 cells and primary human keratinocytes (PHK), respectively. PHKs were isolated from neonatal foreskins following circumcision according to IRB-approved protocols at UAB. They were grown in keratinocyte serum-free medium (Life Technologies) with mitomycin C-treated J2 feeder cells (Swiss 3T3 J2 fibroblasts; Elaine Fuchs, Rockefeller University) [11, 12]. HEK 293 cells were maintained under standard conditions.

### Cytotoxicity and Quantification of Viruses

Antiviral assays: Each experiment that evaluated the antiviral activity of the compounds included both positive (S1 and S2 Figs.) and negative control compounds (data not shown) to ensure the performance of each assay.

Cytotoxicity assays: Concurrent assessment of cytotoxicity was also performed for each study in the same cell line and with the same compound exposure.

All primer/probe sequences used in qPCR and DNA hybridization assays are available in S1 Table.

**S1 Table.**
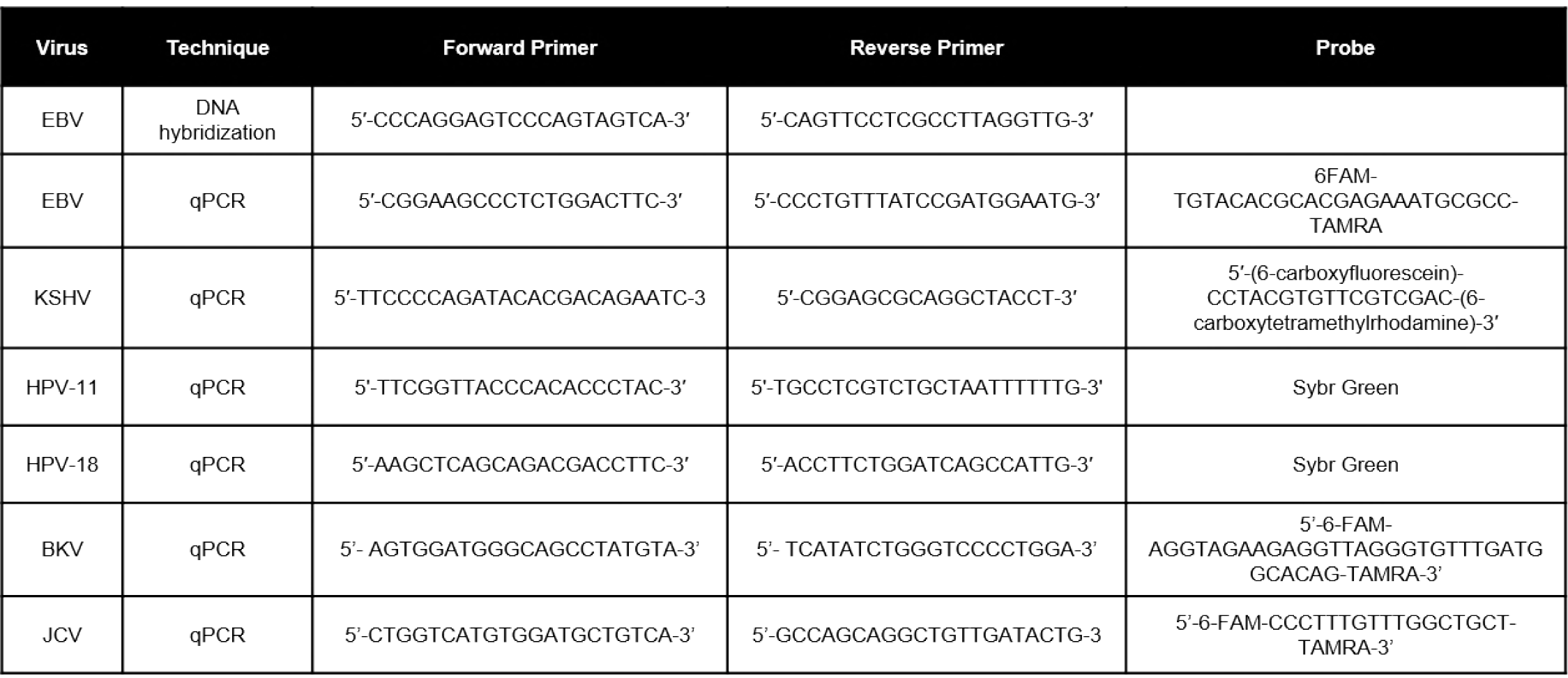
Primers and probes used for qPCR and DNA hybridization

### HCMV

For HCMV cytotoxicity assay was determined via neutral red uptake assay and was conducted as reported previously in HFF cells [13]. Optical densities were determined at 550nm [9].

Yield reduction assays for HCMV were performed on monolayers of HFF cells prepared in 96-well plates and incubated at 37°C for 1d to allow the cells to reach confluency. Media was then aspirated from the wells were infected at a high multiplicity of infection (MOI). At 1h following infection, the inocula were removed and the monolayers rinsed with fresh media. Compounds were then diluted in assay media consisting of MEM with Earl’s salts supplemented with 2% FBS, L-glutamine, penicillin, and gentamycin. Solutions ranging from 0.0032-10µM in primary assay and 0.032-100µM in secondary assay were added to the wells and the plates were incubated for various times, depending on the virus used and represents the length of a single replication cycle for the virus. A duplicate set of dilutions were also performed but remained uninfected to serve as a cytotoxicity control and received equal compound exposure. Supernatants from each of the infected wells were subsequently titered in a TCID_50_ assay to quantify the progeny virus. For the cytotoxicity controls, cytotoxicity was assed using CellTiter-Glo according to the manufacturer’s suggested protocol. For all assays, the concentration of compound that reduced virus titer by 50% (EC_50_) was interpolated from the experimental data.

### MCMV/GPCMV

Cytotoxicity assay was performed in a manner analogous to HCMV (see above).

Plaque reduction assays for MCMV, GPCMV were performed on monolayers of GPL (GPCMV) or MEF (MCMV) cells prepared in six-well plates and incubated at 37°C for 2d to allow the cells to reach confluency. Media was then aspirated from the wells and 0.2ml of virus was added to each of three wells to yield 20-30 plaques in each well. The virus was allowed to adsorb to the cells for 1h and the plates were agitated every 15min. Compounds were diluted in assay media consisting of MEM with Earl’s salts supplemented with 2% FBS, L-glutamine, penicillin, and gentamycin. Solutions ranging from 0.0032-10µM (primary) or 0.032-100µM (secondary) were added to triplicate wells and the plates were incubated for various times, depending on the virus used. For MCMV and GPCMV, the cell monolayer was stained with 1% neutral red solution for 4h at which time the stain was aspirated and cells were washed with PBS. For all assays, plaques were enumerated using a stereomicroscope and the concentration of compound that reduced plaque formation by 50% (EC_50_) was interpolated from the experimental data.

### EBV

Cytotoxicity assay was performed in a manner analogous to KSHV (see above).

For Primary assay, Akata cells latently infected with EBV were induced to undergo lytic infection by addition of 50μg/ml of a goat anti-human IgG [14] and a dose-range of verdinexor (0.032 to 100 μM) was added for 72hr. Denaturation buffer was added and an aliquot aspirated through an Immobilon nylon membrane (Millipore, Bedford, MA). Dried membranes were equilibrated in DIG Easy Hyb solution (Roche Diagnostics, Indianapolis, IN) at 56°C. Specific DIG-labeled probes for EBV (S1 Table). (Roche Diagnostics). Membranes with EBV DNA were hybridized overnight followed by washes in 0.2× SSC and 0.1× SSC both with 0.1% SDS. Detection of specifically bound DIG probe was performed with anti-DIG antibody (Roche Diagnostics). An image of the photographic film was captured and quantified with QuantityOne software (Bio-Rad), and the EC_50_ were interpolated [15].

A plasmid containing the amplified region was diluted to produce the standards used to calculate genome equivalents. Samples were run in duplicate and copy number was calculated.

Secondary assay for EBV were performed in Akata cells that were induced to undergo a lytic infection with 50µg/ml of a goat anti-human IgG antibody by methods we reported previously [16]. Experimental compounds were diluted in round bottom 96-well plates to yield concentrations ranging from 0.0032-100µM. Akata cells were added to the plates at a concentration of 4×10^4^cells per well and incubated for 72h. Assay plates were incubated for seven days at 37°C. For all assays, the replication of the virus was assessed by the quantification of viral DNA. For primer sequences, please see S1 Table. EBV DNA was purified according to manufacturer’s instructions (Wizard SV 96 Genomic DNA Purification System). The purified DNA was then subjected to qPCR that amplified a fragment corresponding to nucleotides 96802-97234 in the EBV genome (AJ507799). Plasmid pMP218 containing a DNA sequences corresponding to nucleotides 14120-14182 (AF148805.2) was used to provide absolute quantification of viral DNA. Compound concentrations sufficient to reduce genome copy number by 50% were calculated from experimental data.

### KSHV

For KSHV, cell viability was assessed with the CellTiter-Glo Luminescent Cell Viability Assay (Promega) using manufacturer’s protocol. Luminescence was quantified on a luminometer. Standard methods were used to calculate CC_50_ [15].

To quantify virus, compound was diluted from 0.008-100µM and added to BCBL-1 cells induced to undergo a lytic infection by the addition of 100ng/ml phorbol 12-myristate 13-acetate (PMA) (Promega). After 7d, total DNA was prepared, and viral DNA was quantified by qPCR (S1 Table for primer/probe sequences). Plasmid pMP218 containing a DNA sequence corresponding to nucleotides 14120-14182 (AF148805.2) was used to provide absolute quantification of viral DNA and EC_50_ [10].

#### AdV

For AdV cytotoxicity assay, at 6d post-infection plates were stained with tetrazolium-based MTS (CellTiter96 Reagent, Promega) and the microtiter plates were read spectrophotometrically at 490/650nm (Molecular Devices).

Prior to assessing cell viability using MTS, supernatants were collected from plates and titrated to determine virus yield. Serial dilutions of supernatants were incubated on HeLa cells for 3 days. Following incubation, wells were visually scored for infection and TCID_50_, EC_50_, CC_50_ and SI were calculated for each sample.

### BKPyV

Cytotoxicity assay was performed in a manner analogous to KSHV above.

Primary assays for BKPyV were performed in 384-well plates containing monolayers of HFF cells. Compound dilutions ranging from 0.0032-100µM were prepared in plates containing cells which were subsequently infected at an MOI of 0.001. After a 7d incubation, total DNA was prepared using Extracta and genome copy number was quantified by qPCR using the primers and probe listed in S1 Table [17]. Plasmid pMP526 served as the DNA standard for quantification purposes. Compounds were confirmed in a similar assay in 96-well plates according to established laboratory protocols with the compounds added 1h post-infection to identify compounds that inhibit early stages of replication including adsorption and entry. Genome copy number was determined by methods described above.

### JCPyV

Cytotoxicity assay was performed in a manner analogous to KSHV above. Evaluation of compounds against JCPyV virus was done in COS7 cells [18] with the plasmid pMP508 to provide a standard curve for quantification.

Primary evaluation of compounds against JC polyoma virus were also performed by methods similar to those for BK virus primary assays but were done in COS7 cells and utilized the MAD-4 strain of JCV. Viral DNA was quantified (S1 Table) together with the plasmid pMP508 to provide a standard curve for absolute quantification. Secondary assays against JCV were also performed in COS7 cells by methods similar to those for BK virus to identify if verdinexor inhibited adsorption or entry of the virus.

### HPV

HPV cytotoxicity assay was performed at the time of viral harvest by scoring cells in a BioRad Automatic Cell Counter for cell viability.

Quantification of virus was determined via transient replication of an HPV-11 replication origin-containing plasmid in HEK293 cells co-transfected with an HPV-11 replicative DNA helicase E1 and origin binding E2 protein [19]. 293 cells were cultured with test compounds added 4hr post-transfection for 2 days. Low molecular weight DNA was isolated, digested with *Dpn1* and exonuclease III to eliminate unreplicated DNA. The *Dpn1*-resistant DNA was then subjected to qPCR analyses amplifying a small portion of the replication origin [19]. In a negative control, the E1 expression vector was omitted or cultures were treated with the inhibitor cidofovir.

For secondary assays, the amplification of HPV-18 DNA was determined in an organotypic squamous epithelial raft culture of PHKs. Whole genomic HPV-18 plasmids were generated in plasmid-transfected PHKs and developed into organotypic cultures as described [11]. Based on the results of a 3-day toxicity assay, concentrations of test compound were added to a subset of raft cultures from day 6 through day 13 when the cultures were harvested. Similarly treated normal PHK raft cultures were also prepared to assess toxicity of uninfected tissue. One set of cultures were fixed in 10% buffered formalin, paraffin embedded and sectioned for in situ assay. Sections were stained with hematoxylin and eosin. Total DNA harvest from unfixed raft cultures was used to determine the HPV-18 copy number/per cell by qPCR. Inhibition was expressed as % of viral DNA copies relative to the untreated cultures.

## Results

### EBV replication is antagonized by treatment with verdinexor

In this study, we determined the efficacy of verdinexor to inhibit replication *in vitro* against EBV infection of Akata cells, with EC_50_ values in the range of >0.48µM (primary screen) to 0.05µM (secondary screen) (Fig. 3). This correlated to an SI of <1 in primary screen measured by DNA hybridization and 7 in a secondary assay that used qPCR. Interestingly, a similar pattern was observed in control-treated cells, with acyclovir administration resulting in EC_50_ values of 15.03µM in primary assay and 7.90µM in a secondary screen (S1 Fig.). This correlated to SI values of >7 and >13, respectively. These results indicate that inhibition of nuclear export with verdinexor is effective in treating EBV infections at nanomolar concentrations when measured by DNA hybridization.

**Figure 3.**
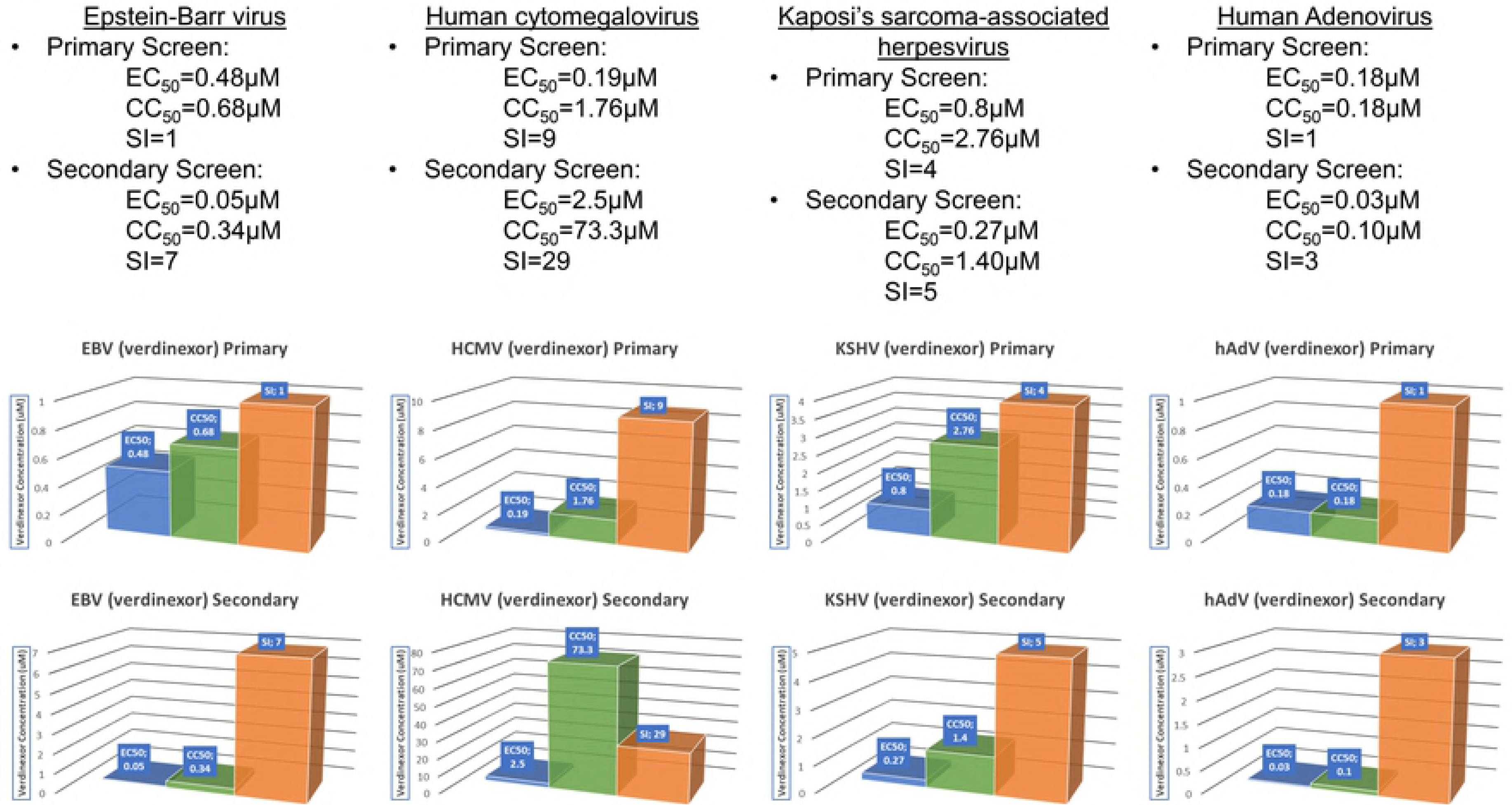
In vitro antiviral and cytotoxicity levels for verdinexor. Results of verdinexor treatment against viral infections. EC_50_ values are plotted in blue, CC_50_ values are plotted in green, and the SI value for each assay is plotted in orange.

### Verdinexor treatment affects viral replication of HCMV in HFFs

Verdinexor inhibition of HCMV replication was potent in both primary and secondary screens, with EC_50_ values of 0.19µM and 2.5µM, respectively (Fig. 3). Taking cytotoxicity into consideration, we observed an SI of 9 in the primary screen, and 29 in the secondary screen. Treatment of HCMV-infected HFF cells with the nucleoside analogue ganciclovir resulted in EC_50_ values of 0.51µM in primary screen and 4µM in secondary with SI values of >196 and >25 respectively (S1 Fig.). With these data in hand, we were next interested if replication of cytomegaloviruses tropic for small animals would be efficiently inhibited by verdinexor treatment.

### Verdinexor is efficacious in lowering replication of MCMV in MEF cells and GPCMV in GP Lung cells

Verdinexor treatment of murine CMV and guinea pig CMV infections was highly efficacious in both mouse and guinea pig infected cell lines, with EC_50_ values of 0.19µM and SI values of >789 and >11, respectively (Fig. 4). These data, along with results from the HCMV screen indicate verdinexor possesses a potentially favorable pharmological profile to treat CMV infection, and warrants further assessment in small animal models of CMV infection such as mice and guinea pigs.

**Figure 4.**
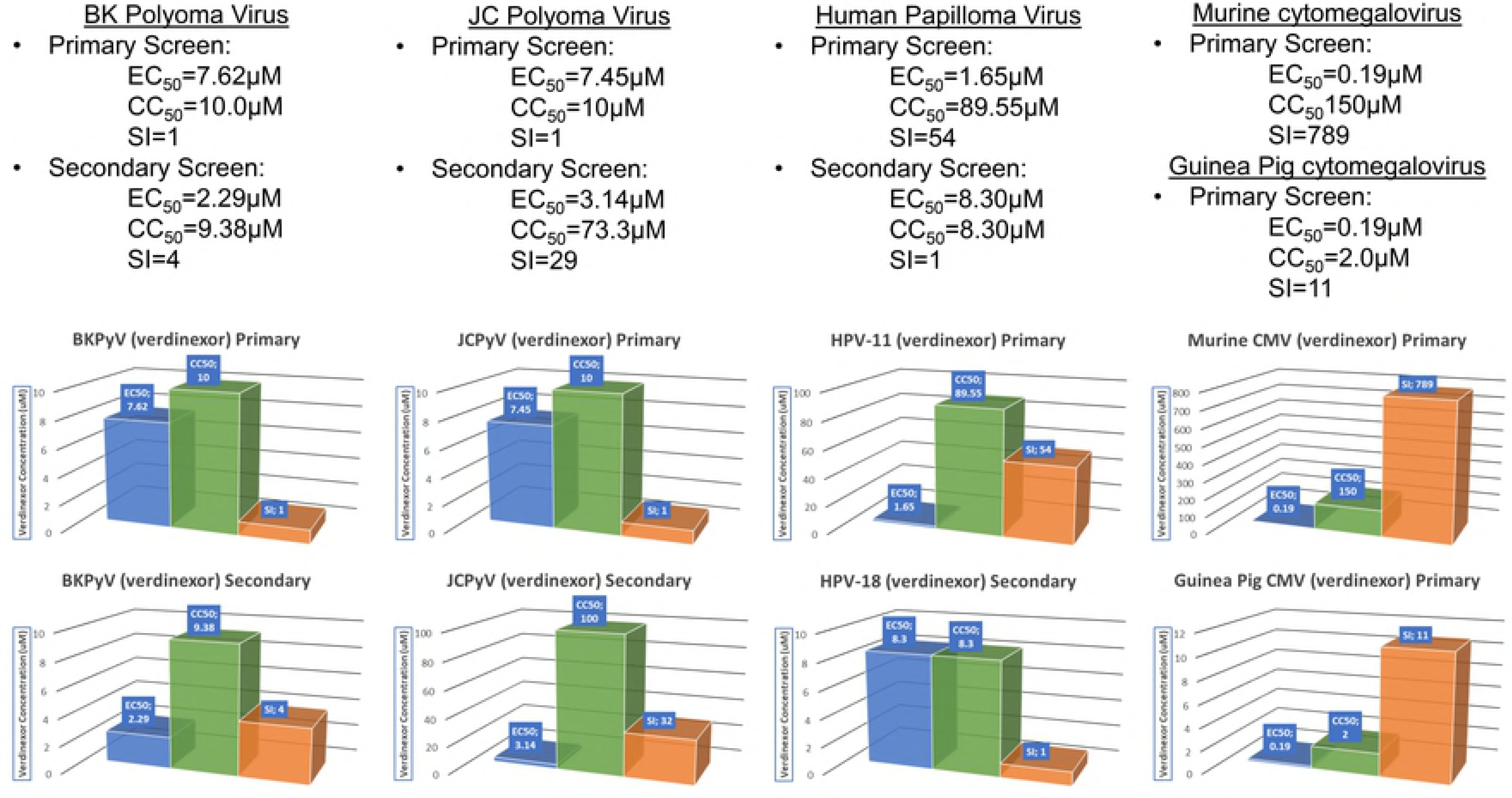
In vitro antiviral and cytotoxicity levels for verdinexor. Results of verdinexor treatment against viral infections. EC_50_ values are plotted in blue, CC_50_ values are plotted in green, and the SI value for each assay is plotted in orange.

### Verdinexor is efficacious in an in vitro model of KSHV infection

We observed verdinexor inhibition of KSHV replication at EC_50_ concentrations at or below those for cidofovir, a viral DNA polymerase inhibitor used as a positive control. In BCBL-1 cells, primary screening with verdinexor produced an EC_50_ value of <0.8µM and a SI of >3.45 (Fig. 3). This compares to cidofovir, which was observed to have an EC_50_ of less than 0.48µM however, due to the low toxicity observed in BCBL-1 cells had an SI of >125 (S1 Fig.). As per protocol, we next performed a secondary screen analogous to primary screening. This time, verdinexor demonstrated 50% inhibition at a concentration of 0.27µM resulting in a SI of 5.2 (Fig. 2), while cidofovir produced an EC_50_ of 1.68µM with a concomitant SI of >36 (S1 Fig.). Despite differing CC_50_ values between the screens, these results provide rationale for further examination of the efficacy of verdinexor against KSHV in models of KSHV infection. Verdinexor also has anti-oncogenic properties, opening the possibility of an alternative mechanism of inhibiting KSHV-induced malignancies.

### Adenovirus infection is inhibited by treatment with verdinexor

Initial testing of verdinexor against Ad5 infection in HeLa cells demonstrated efficacy with an EC_50_ value of >0.18µM (Fig. 3). However, these cells proved to be highly sensitive to the cytotoxic effects of antiviral treatment and as such we did not observe efficacious SI values. In a secondary screening, the EC_50_ value for verdinexor was <0.03µM and an SI of >3 (Fig. 3). A highly potent neutralizing anti-Ad5 monoclonal antibody was used as a positive control, and treatment resulted in SI of >50 and >41 in primary and secondary assays, respectively (S1 Fig.). With this *in vitro* data, we feel that further studies of verdinexor efficacy against AdV infection should be considered using different non-cancerous cell lines to address observed cytotoxicity.

### Efficacy of verdinexor against the polyomaviruses BKPyV and JCPyV

Treatment with verdinexor against BKPyV infection in HFF cells resulted in an SI of >1 in primary assays, and 4 in secondary screen with EC_50_ values of 7.62 and 2.29µM respectively when measured by qPCR (Fig. 4). Cidofovir treatment resulted in EC_50_ values of 4.48 and 2.06µM, with concomitant SI of >22 and >49 (S2 Fig.). JCPyV virus was observed to have EC_50_ of 7.45µM with a SI >1 in COS-7 cells. Administration of cidofovir resulted in an EC_50_ of 3.86µM and an SI of 17. Thus, despite similar efficacious concentrations of drug, cytotoxic levels varied between drugs in these cells resulting in differences in SI. The results of additional suggest additional testing of BKPyV and JCPyV in alternate cell lines will serve to assess the true ability of verdinexor to inhibit polyomavirus replication.

### Efficacy of verdinexor against 2 strains of HPV in 2 different cell lines

In a primary screen of verdinexor we observed extremely potent inhibition of HPV-11 replication, with an SI of 54 and EC_50_ value of 1.65µM when assayed in HEK293 cells (Fig. 4). This compared well with cidofovir treatment, which produced an EC_50_ of 148µM and a SI of only >1 (S2 Fig.). In a secondary screen using PHKs and HPV-18, verdinexor did not show efficacy, due in large part to the high degree of cytotoxicity observed in these cells. Interestingly we observed similar lack of efficacy in the control treatment cells, with the MEK inhibitor Uo_126_ producing an SI of >1 (S2 Fig.). This appears to be either a virus strain or cell type-specific result, and may not be indicative of verdinexor and/or Uo_126_ performance *in* vivo, especially in light of the results of the primary screen with HPV-11. Taken together, these data suggest verdinexor may have the potential to serve as a type-specific and potent antiviral compound for the treatment of certain HPV-related diseases.

## Discussion

Opportunistic infections associated with immunodeficiency are a burgeoning field of translational research. Limited therapies exist to treat these normally benign infections, and with increasing numbers of patients displaying symptoms of immunodeficiency, the need for novel strategies by which to treat opportunistic viral infections is high.

Verdinexor is a member of a class of small molecules which bind the nuclear export protein XPO1 to inhibit its function. The result is nuclear accumulation of proteins (nearly 220 identified) that utilize XPO1 for translocation into the cytoplasm. Verdinexor has shown promising potential as a broad-spectrum antiviral drug [6, 7, 20]. It demonstrates a dual mechanism of action, by inhibiting replication of viruses that utilize XPO1 machinery for replication, and relieving virus-induced inflammation [6]. Importantly, inhibition of XPO1 does not rely on immune status for its antiviral effects, and viral resistance is minimized by targeting a host cell protein. In a study with influenza A, 10 passages of the virus in the presence of verdinexor resulted in highly attenuated resistant mutants with altered in vitro growth kinetics [21]. We believe verdinexor offers a novel therapy that targets the underlying causes of immunopathology (namely inflammation) in immunocompromised individuals while simultaneously inhibiting viral replication that can occur when opportunistic viruses develop active infection in these patients.

Verdinexor was effective in inhibiting EBV replication in Akata cells, with EC_50_ as low as 50nM and SI of 7 when measured by qPCR. Acyclovir was observed to have higher EC_50_ and lower SI values when measured by DNA hybridization, a pattern also observed for verdinexor treatment. This may be an indication that qPCR was more sensitive for measuring subtle differences in viral titer. We hypothesize that the efficacy of verdinexor could be explained by the dependence of the EBV protein SM on XPO1-mediated nuclear export (Fig. 5) [22]. SM is an adaptor protein involved in the nucleocytoplasmic export of mRNAs encoding lytic EBV genes. Blockade of this export should prevent shuttling of these mRNAs to the cytoplasm for translation. Thus, this proposed mechanism of action, inhibition of viral translation, may be more accurately measured by qPCR.

**Figure 5.**
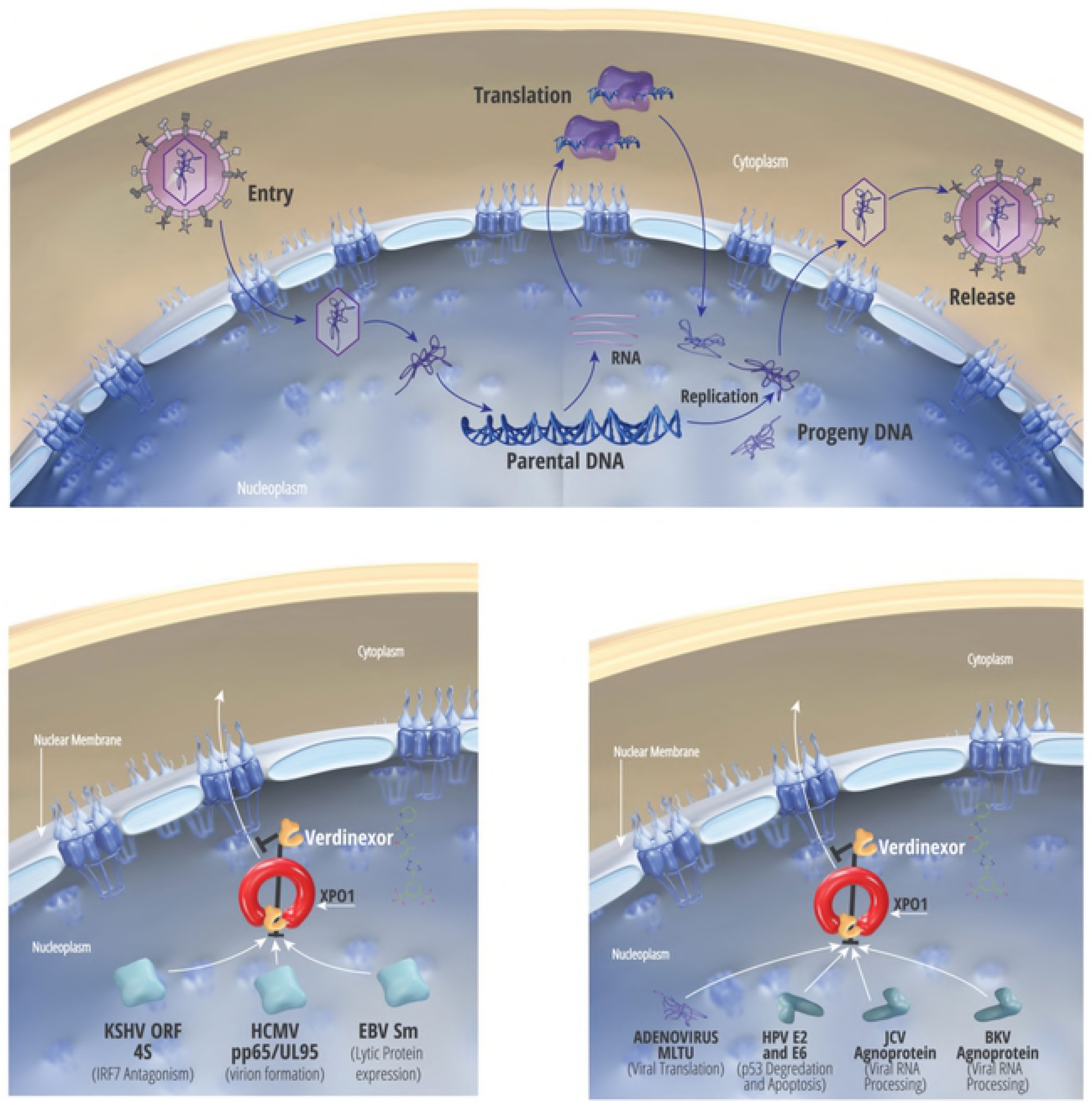
Proposed model of antiviral activity of verdinexor against opportunistic dsDNA viruses. Likely mechanism of action of verdinexor’s antiviral effects on each of the viruses tested. Model is based on data in the literature.

We observed a SI value for verdinexor treatment of HCMV-infected cells exceeding that of ganciclovir positive control. One possible explanation from the literature is the requirement of HCMV tegument protein pp65 (UL83) to localize to the cytoplasm where it participates in virion assembly (Fig. 5). Prior to its cytoplasmic localization, pp65 is found in the nucleus where it utilizes XPO1 for its export [23]. Blockade of XPO1 by verdinexor presumably causes accumulation of pp65 in the nucleus, where it is unable to participate in virion assembly. Nuclear sequestration of another tegument protein, UL94, a protein carrying an XPO1-dependent NES sequence would be predicted to hinder viral replication as well. We also observed very promising SI values for verdinexor treatment of murine CMV and guinea pig CMV-infected cells, warranting further investigation of verdinexor treatment in *in vivo* mouse and guinea pig models.

In KSHV, blockade of XPO1-dependent nuclear export with verdinexor was shown to reduce viral replication, likely via inhibition of multifunctional viral protein ORF45 (Fig. 5). ORF45 is normally localized to the cytoplasm and inhibits IRF-7 translocation to the nucleus [24]. Nucleocytoplasmic trafficking of ORF45 occurs in an XPO1-dependent manner [25], and blockade of this export by verdinexor is speculated to sequester ORF45 to the nucleus. Interestingly, although verdinexor demonstrated comparable EC_50_ values to those of cidofovir, toxicity values in the BCBL-1 cells varied significantly, resulting in higher SI values for cidofovir despite similar EC_50_ concentrations. The qPCR assay used to determine EC_50_ values could be more sensitive to changes in genome replication such as those conferred by treatment with cidofovir, whereas verdinexor is expected to affect viral assembly and host immune responses. Additionally, the oncogenic nature of BCBL-1 cells would be expected to be highly sensitive to the anti-oncogenic properties of verdinexor (see below).

Adenoviruses are some of the most ubiquitous viruses in the human population. We observed verdinexor inhibition of viral replication at concentrations <30nM. Adenoviral protein transcription is thought to be dependent on XPO1-mediated nuclear export of the viral major late transcription unit (MLTU; Fig. 5), which is critical for replication of the virus in the cytoplasm [26]. Inhibition of this nuclear export by verdinexor should have a profound effect on viral titers. We did not observe high SI values in either screen with verdinexor, perhaps due to the general anti-oncologic effect of SINE compounds on the cells utilized in the assay. Inhibition of XPO1 results in sequestration of inflammatory mediators and tumor suppressor genes in the nucleus of oncogenic cells, where they exert their effects. HeLa cells are a well-known cancerous cell line and it would be predicted that verdinexor would naturally have detrimental effects on the survival of HeLa cells. Thus, assays in different, preferably primary, cells would be prudent.

BKPyV virus can cause neuropathy and malignancies [27] while JCPyV causes leukemias, lymphomas, and progressive multifocal leukoencephalopathy (PML), a rare but routinely fatal manifestation that leads to the destruction of myelin and oligodendrocytes in the central nervous system, incidence of which has increased due to use of immune altering biologic treatments for autoimmune disorders. Polyomavirus treatment with verdinexor resulted in EC_50_ values nearly equivalent to those observed for cidofovir treatment. The agnoprotein of these viruses utilize XPO1 for nucleocytoplasmic shuttling [28]. Sequestration of agnoprotein to the nucleus (Fig. 5) of infected cells likely has effects on viral replication, as it is posited to be involved in viral transcription and replication [29–31], virion assembly [32, 33], and DNA encapsidation [34]. Agnoprotein can halt cell cycle progression in the G2/M phase [35], inhibit DNA repair mechanisms [36], and importantly for PML pathogenesis induce apoptosis in oligodendrocytes [37]. Thus, on the basis of current knowledge, we believe there are multiple mechanisms of viral inhibition of polyomaviruses by verdinexor.

Verdinexor inhibition of HPV replication in a primary screen using HPV-11 in HEK293 cells produced an SI of 54. This compared favorably to cidofovir treatment which interestingly did not show efficacy against this virus when assayed by qPCR. A secondary screen using HPV-18 in PHKs found that PHK cells are more sensitive to drug-induced cytotoxicity, however this appears to be cell type-specific. It is speculated that the observed results in the primary screen may be a result of HPV utilizing XPO1 for a number of pathogenic mechanisms (Fig. 5). In HPV-positive cancer cells, the HPV E6-dependent pathway of p53 degradation is active and required for growth. Inhibition of XPO1 in HPV-positive tumor cells should result in nuclear accumulation of intact p53, suggesting that E6-mediated degradation of p53 is dependent on its nuclear export [38]. Additionally, XPO1-mediated nuclear export of the viral protein E2 is responsible for cellular apoptosis in high-risk HPV genotypes [39], so nuclear sequestration of this protein may also play a role in verdinexor’s inhibitory properties. It is important to note that there exists an extremely effective vaccine to prevent HPV-11 and −18 diseases. Efficacy, however, wanes with age of administration, as older individuals have more likely been exposed to HPV than adolescents (for whom the vaccine is indicated for). As such, there remains a large population the will not benefit from vaccination, and antiviral drugs will be needed to combat their infections.

The results of this in vitro screen of verdinexor against opportunistic viruses of the immunocompromised resulted in a wide range of results. Indeed, due to variability in antiviral assays performed at different times, we chose to perform secondary assays on all human viruses tested to gain greater confidence in our observed results. In nearly every case, SI values in the secondary screen were higher than in the primary screen, reflecting perhaps increased experimental acumen, differential assays used to measure cytotoxicity and viral inhibition, or a plethora of other uncontrollable variables. Importantly, verdinexor possesses potent myelosuppressive effects that makes the compound innately toxic to oncogenic cell lines [40], as would be expected since its analogous compound, selinexor, is currently in advanced human trials for multiple malignancies. This anti-oncogenic property of verdinexor, a result of sequestration of tumor suppressor proteins and pro-apoptotic factors in the nucleus of cancerous cells, is toxic to most immortalized cell lines and results in a very low concentration to reach 50% cytotoxicity. This in turn results in a lowered SI value, and is likely the reason we observed reduced SI values (ie. below 10) for many of the viruses tested. This can be easily observed in the assays for EBV, which were performed in the cancerous Akata cell line that latently expresses EBV (which itself is oncogenic) and resulted in CC_50_ values below 680nM. This is in stark cocontrast to HCMV, assays performed in primary HFF cells and resulting in relatively high CC_50_ concentrations (1.76µM in primary screen and 73.3µM in secondary) and concomitant increased SI values of 9 and 29, respectively. Thus the cytotoxicity observed in some of these experiments may not be indicative of verdinexor behavior in vivo, and that long-term application of verdinexor may not be as toxic as these in vitro assays may indicate. Indeed, verdinexor’s analogous anti-oncology drug selinexor has been administered to some patients for over 2 years at higher doses than utilized in this study when calculated for human administration. As such, cytotoxicity in assays using oncogenic cell lines should be interpreted cautiously, and further studies with these viruses in primary cell lines or in vivo should be conducted where possible.

Individuals with weakened immune responses face severe disease from benign virus infections. Treating these diseases can be challenging due to impaired immune function. Thus, it is imperative to find novel antiviral therapies that target host or viral proteins and are able to work independently of the immune system. Verdinexor presents a promising approach to target both HIV and opportunistic viruses. As demonstrated by the broad antiviral screening data, verdinexor demonstrates efficacy against a diverse panel of viruses with a median EC_50_ value of 2μM (range 30nM-8.3μM) and SI ranging from 1-789. Treatment of cells with verdinexor sequesters XPO1-cargo proteins in the nucleus, including (presumably) a wide variety of viral proteins (Figs. 2 and 3). In addition to sequestering viral proteins and RNA to the nucleus, XPO1 inhibition leads to nuclear retention of host factors essential to viral replication such as enzymes and initiation factors, and inflammatory host factors such as cytokine, mRNA and transcription factors responsible for immunopathology in the infected individual. These data, along with our previous findings from HIV studies, justify a future evaluation of verdinexor in models of multiple viral infections.

## Acknowledgements

We wish to acknowledge the laboratories of Dr. Mark N. Prichard (UAB), Dr. Louise T. Chow (UAB), and Dr. Roger Ptak (Southern Research Institute) for performing the contracted in vitro screens. We thank Susie Harrington for assistance in compiling figures and T.J. Unger and Lori King for critical review of the manuscript.

## Supporting Information

**S1 Fig. In vitro antiviral and cytotoxicity levels for positive control compounds.**

Results of positive control treatment against viral infections. EC_50_ values are plotted in blue, CC_50_ values are plotted in green, and the SI value for each assay is plotted in orange.

**S2 Fig. In vitro antiviral and cytotoxicity levels for positive control compounds.**

Results of positive control treatment against viral infections. EC_50_ values are plotted in blue, CC_50_ values are plotted in green, and the SI value for each assay is plotted in orange.

